# Targeting Mesothelin via the native T cell receptor

**DOI:** 10.64898/2025.12.02.689353

**Authors:** Kiriakos Koukoulias, Ryu Yanagisawa, Alejandro G. Torres Chavez, Penelope G. Papayanni, Yovana Velazquez, Spyridoula Vasileiou, Ann M. Leen

**Affiliations:** Center for Cell and Gene Therapy, Baylor College of Medicine, Texas Children’s Hospital, and the Methodist Hospital, Houston, Texas, USA

## Abstract

Mesothelin (MSLN) is a GPI-anchored cell surface glycoprotein that is overexpressed in various solid tumors, including mesothelioma, triple-negative breast cancer, colon, ovarian and pancreatic cancer, with restricted normal tissue expression. To explore the immunogenicity and immunotherapeutic potential of MSLN to T cells with native receptor specificity, 29 individuals of diverse HLA backgrounds were interrogated for T cell activity against MSLN. Twenty one (72%) subjects (21/29) mounted a specific T cell response when repetitively challenged with MSLN antigen. Reactive cells were Th1 polarized, polyfunctional, predominantly detected in the CD8^+^ T cell compartment and cytotoxic toward autologous and MSLN^+^/HLA-matched tumor cell lines in conventional 2D in vitro assays. Furthermore, these cells produced potent anti-tumor effects in a novel 3D tumor spheroid model system established to evaluate the safety and potency of reactive cells against tumors including pancreatic, cervical, and colorectal cancer and mesothelioma. Taken together, these findings establish the feasibility of targeting MSLN using adoptively transferred T cells with native antigen specificity.

## Introduction

Mesothelin (MSLN) is a cell surface glycoprotein expressed by mesothelial cells lining the pleura, peritoneum, and pericardium. Initially synthesized as a 69kDa precursor protein, MSLN is cleaved post-translationally resulting in a 40kDa membrane-bound C-terminal fragment and a 31kDa N-terminal secreted fragment [also known as the megakaryocyte potentiating factor (hMPF)]^1, 2, 3, 4,5^. In humans the physiologic role of MSLN remains unclear but mice with homozygous null mutations show no detectable anatomic, developmental or reproductive defects^6^. However, MSLN is overexpressed in many tumors, including mesothelioma, pancreatic, ovarian, colon, uterine, cervical and triple negative breast cancer (TNBC)^7, 8, 9, 10, 11, 12, 13^, and in the malignant setting there have been reports highlighting a potential role in cancer cell adherence, survival, proliferation, tumor progression and chemoresistance^8, 9, 14, 15, 16, 17, 18, 19, 20, 21^.

As MSLN is overexpressed on the surface of many tumors and has minimal normal tissue expression, it has been investigated extensively as a target for antibody-based therapeutics including recombinant immunotoxins and monoclonal antibodies^5, 22, 23, 24, 25^, as well as chimeric antigen receptor (CAR) T cells^26, 27, 28, 29, 30, 31, 32^. In the current study, however, we sought to explore MSLN as a target for T cells via the native T cell receptor, which recognizes antigen that is processed and presented by human leukocyte antigen (HLA) molecules. We reasoned that such an approach would produce potent and durable therapeutic benefit and be impervious to the impact of antigen cleavage from the tumor cell surface.

We now examine the immunogenicity of MSLN to endogenous T cells isolated from individuals of diverse HLA backgrounds and investigate the effector profile and immunotherapeutic potential of antigen-specific CD8^+^ and CD4^+^ T cells. Here we report on the immunodominance, phenotypic characteristics, and functional activity of reactive cells and map an array of novel, HLA-restricted CD4^+^ and CD8^+^ T cell peptide epitopes. We also interrogate the capacity of reactive cells to mediate potent antitumor effects in multiple 2D and 3D models of MSLN-expressing cancers.

## Materials and Methods

### Donors and cell lines

Peripheral blood mononuclear cells (PBMCs) [obtained from healthy adult volunteers with informed consent using Baylor College of Medicine institutional review board-approved protocol (H-45017)] were used to generate dendritic cells (DCs), MSLN-specific T cells (MSLN-STs), and phytohaemagglutinin (PHA) blasts. Donors were selected based on (i) HLA typing and (ii) availability for repeated blood draws. Sex/gender considerations were not taken into account for donor selection. PHA blasts were generated from PBMC (0.5 × 10^6^/ml) using 5 µg/ml PHA (Sigma-Aldrich, St. Louis, MO) and cultured in T cell medium [45% RPMI 1640 (HyClone Laboratories, Logan, Utah), 45% Click’s medium (Irvine Scientific, Santa Ana, California), 2 mM GlutaMAX TM-I (Life Technologies, Grand Island, New York), and 10% human AB serum (Valley Biomedical, Winchester, Virginia)] supplemented with 100U/ml interleukin 2 [IL2; Proleukin® (aldesleukin), Yardley, PA], which was replenished every 2 days. A panel of human cancer cell lines, including pancreatic cancer (CFPAC-1, CAPAN-1), cervical carcinoma (MS751), colon adenocarcinoma (SK-CO-1), breast cancer (Hs578T, HCC1806), uterine adenocarcinoma (KLE), ovarian cancer (OVCAR3), prostate cancer (LNCaP), and mesothelioma (NCI-H2052, NCI-H2452, NCI-H28) were purchased from the American Type Culture Collection (ATCC, Manassas, VA). CFPAC-1 and CAPAN-1 cells were cultured in IMDM (Gibco, Waltham, MA); MS751 and SK-CO-1 in EMEM (ATCC); Hs578T and KLE in DMEM (HyClone Laboratories), with KLE additionally supplemented with Ham’s F-12 (Gibco); and OVCAR3, HCC1806, LNCaP, NCI-H2052, NCI-H2452, and NCI-H28 in RPMI-1640. All media were supplemented with 10% fetal bovine serum (FBS) (Gibco). Cells were cultured at 37 °C in a humidified atmosphere containing 5% CO₂.

### MSLN Peptides/pepmixes

For antigen-specific stimulation, we used peptide mixtures (pepmixes) (15-mers overlapping by 11 amino acids) spanning MSLN, which were purchased from Genemed Synthesis, Inc. (San Antonio, TX) and LifeTein, LLC (Somerset, NJ). For epitope mapping, peptides were pooled into 25 mini-pools, each containing 12 or 13 15-mer peptides, organized such that each peptide was uniquely present in two mini-pools. To identify minimal epitopes, we generated a series of shorter peptides ranging from 14 to 9 amino acids in length, spanning immunogenic sequences of MSLN and covering all potential peptide epitopes. Lyophilized peptides were reconstituted at 10 mg/mL in dimethyl sulfoxide (DMSO (Sigma-Aldrich).

### MSLN-STs generation

#### DC generation

Autologous DCs were used as antigen-presenting cells (APCs) and generated by isolating monocytes from PBMCs by CD14 selection using MACS Beads (Miltenyi Biotec, Bergisch Gladbach, GmbH). Subsequently the cells were cultured in DC medium (CellGenix, Freiburg, GmbH) with 800 U/ml granulocyte macrophage-colony stimulating factor (GM-CSF) (Sargramostim Leukine; Immunex, Seattle, WA) and 400 U/ml IL-4 (R&D Systems, Minneapolis, MN) for 5 days. IL-4 and GM-CSF were replenished on day 3 and on day 5, DCs were matured in DC medium supplemented with 10 ng/ml IL-1β (R&D Systems),10 ng/ml tumor necrosis factor (TNF)-α (R&D Systems), 1 µg/ml prostaglandin E2 (Sigma-Aldrich), 10ng/ml IL6 (R&D Systems), 800 U/ml GM-CSF and 400 U/ml IL-4.

#### MSLN T cell generation

On day 6 or 7 of culture, mature DCs were pulsed for 30-45 minutes at 37 °C with 200ng of MSLN peptide. CD14-negative PBMCs were used as responder cells and stimulated with MSLN pepmix-pulsed DCs at a stimulator:responder (S:R) ratio of 1:10. Cells were cultured at 1 × 10⁶/ml in T cell medium supplemented with IL-7 (10 ng/ml), IL-12 (10 ng/ml), IL-15 (5 ng/ml), and IL-6 (10 ng/ml) (R&D Systems). On day 9 or 10, MSLN-STs were harvested, counted using trypan blue exclusion to distinguish live and dead cells, resuspended at 0.5 × 10⁶/ml in T cell medium supplemented with IL-7 and IL-15, and then restimulated with MSLN pepmix-pulsed DCs at a S:R ratio of 1:10. On day 7, cultures were fed with fresh T cell medium supplemented with IL-15. Thereafter, cultures were stimulated weekly with pepmix-pulsed DCs at a S:R ratio of 1:10 and fed with fresh T cell medium supplemented with IL-15.

### Immunophenotyping

MSLN-STs were surface-stained with monoclonal antibodies against CD25 (RRID:AB_400334), CD28 (RRID:AB_400368), CD69 (RRID:AB_400054), CD45RO (RRID:AB_400511), CD279 (PD-1) (RRID:AB_2159176), CCR7 (CD197) (RRID:AB_10561679) [Becton Dickinson (BD), Franklin Lakes, NJ], CD3 (RRID:AB_2876783), CD4 (cat# A96417), CD8 (cat# A82791), CD16 (cat# IM1238U), CD56 (RRID:AB_131195), CD62L (cat# IM2713U) (Beckman Coulter, Brea, CA), CD366 (TIM-3) (RRID:AB_2561717), and CD223 (LAG3) (RRID:AB_2629755) (BioLegend, San Diego, CA). A total of 2 × 10⁵ cells were pelleted in phosphate-buffered saline (PBS) (Sigma-Aldrich), followed by the addition of antibodies in saturating amounts (5 µl) and incubation for 15 minutes at 4°C. The cells were then washed, resuspended in 300 µl of PBS, and at least 20,000 live cells were acquired on a Gallios™ flow cytometer (Beckman Coulter). To detect MSLN expression in cancer cell lines, 2 × 10⁵ cells were washed twice and stained with 2 µl of a monoclonal, anti-human, recombinant MSLN antibody (RRID:AB_2733680) (Miltenyi Biotec), diluted 1:50 in PBS containing 0.5% bovine serum albumin (BSA) (Sigma-Aldrich). After 10 minutes of incubation at 4°C in the dark, cells were washed twice, resuspended in 200 µl of PBS containing 0.5% BSA, and approximately 20,000 live cells were acquired using a CytoFLEX flow cytometer (Beckman Coulter). Flow cytometry data were analyzed using Kaluza® software version 2.1 (Beckman Coulter).

### Functional studies

#### Enzyme-linked immunospot assay (ELISpot)

ELISpot analysis was used to quantify the frequency of IFNγ- or Granzyme B (GrB)-secreting cells. Briefly, MSLN-STs were resuspended at 1–2 × 10⁶ cells/ml in T cell medium, and 100 µl of cells was added to each ELISpot well. Antigen-specific activity was measured after direct stimulation with the stimulating peptides. After 16–18 hours of incubation, plates were developed according to manufacturer’s instructions (Mabtech, Inc., Cincinnati, OH), dried overnight at room temperature, and then quantified using the IRIS ELISpot/FluoroSpot reader (Mabtech, Inc.). The breadth of activity in MSLN-STs generated was evaluated by IFNγ ELISpot using the 25 mini-peptide pools, each containing 12 or 13 peptides, representing the entire MSLN sequence. MSLN-STs were stimulated with each of the mini-pools to identify 15-mer peptides that contained immunogenic peptide epitopes. Subsequently, fine mapping studies were performed using a panel of 14–9-mer peptides spanning the sequence of the stimulatory 15-mer peptides. For HLA blocking studies, MSLN-STs were pre-incubated with 10µl of the MHC class I antibody (Santa Cruz Biotechnology, Inc., Santa Cruz, CA) or 10µl DR (Leinco Technologies, Fenton, MO), DQ, DP (Biolegend) antibodies for 30 minutes at 37°C, prior to being added to the ELISpot plate.

#### FluoroSpot

For the quantitation of polyfunctional cells simultaneously secreting IFNγ, GrB and/or TNFα a commercial FluoroSpot assay was used (Human IFNγ/Granzyme B/TNFα FluoroSpot Plus, Mabtech, Inc.). Briefly, MSLN-STs were resuspended at 0.5 to 2×106 cells/ml in T cell medium and 100μl of cells were added to each FluoroSpot well. Antigen-specific activity was measured after direct stimulation with the stimulating peptides. After a minimum of 18 hours of incubation, plates were developed as per manufacturer’s instructions and then visualized and quantified using the IRIS ELISpot/FluoroSpot reader.

#### Intracellular cytokine staining

MSLN-STs were harvested, resuspended in T cell medium (2×10^6^/ml) and 200μl added per well of a 96-well U-bottom plate. Cells were incubated overnight with 200ng of test or control peptides along with Brefeldin A (1μg/ml), monensin (1μg/ml), CD28 and CD49d (1μg/ml) (BD). Next, MSLN-STs were washed with PBS, pelleted, surface-stained with CD8 (cat# IM2469U; Beckman Coulter) and CD3 (RRID:AB_400286) (BD) (5μl/antibody/tube) for 15 minutes at 4°C, then washed, pelleted, fixed and permeabilized with Cytofix/ Cytoperm solution (BD) for 20 minutes at 4°C in the dark. After washing with Perm/Wash Buffer (BD), cells were incubated with 10μl of IFNγ (RRID:AB_400425) and TNFα (RRID:AB_400435) antibodies (BD) for 30 minutes at 4°C in the dark. Cells were then washed twice with Perm/Wash Buffer and at least 50,000 live cells were acquired on a Gallios™ Flow Cytometer and analyzed with Kaluza® Flow Analysis Software.

#### Chromium release assay

We measured cytotoxic activity in 5-8 hour ^51^Cr release assays, using E:T ratios of 40:1, 20:1, 10:1, and 5:1. MSLN-specific T cells were used as effectors and the targets were PHA blasts alone, MSLN pepmix-pulsed PHA blasts or cancer cell lines (CFPAC-1, Capan-1, KLE, MS751, SK-CO-1, NCI-H2452). To identify HLA restriction, we used autologous and partially HLA-matched peptide-pulsed PHA blasts as targets. The percentage of specific lysis was calculated as follows: [(experimental release – spontaneous release)/ (maximum release – spontaneous release)] x 100. For HLA blocking studies, ^51^Cr-labeled targets were pre-incubated with 10µl of the MHC class I or Class II antibodies for 30 minutes at 37°C, prior to adding effector T cells.

#### Luciferase-based cytotoxicity assay using 3D spheroids

Spheroids were generated by seeding 2000 GFP/FFluc^+^ MS751 or CFPAC-1 cells/well in 96-well flat-bottom plates (Corning, NY) that were pre-coated with 45 μL of 1% (w/v) agarose/well (Thermo Fisher Scientific, Pittsburgh, PA). Following spheroid formation (24 hours), partially HLA-matched MSLN-STs or control partially HLA-matched T cells [(Respiratory Syncytial Virus-Specific T cells, (RSV-STs)] were added to the tumor cells at the indicated E:T ratios. Total Green Object Integrated Intensity (expressed as GCU × µm²/image) was monitored using the IncuCyte S3 Live-Cell Analysis System (Sartorius, Gottingen, GmbH) over a 24-hour co-culture period. Quantification was performed following Top-Hat background subtraction analysis.

### Immunohistochemistry

To assess the MSLN expression profile in primary pancreatic tumor samples a tissue microarray (IMGENEX Corporation, San Diego, CA) was placed in an oven at 60°C for 60 minutes, deparaffinized with three changes of xylene for 5 minutes each and then rehydrated through a series of graded alcohols with a final rinse in distilled water. Endogenous peroxides were quenched by soaking sections in 3% methanol hydrogen peroxide for 10 minutes. After washing with distilled water, slides were placed in a steamer for 20 minutes (high pressure) in Target Retrieval solution (Dako Corporation, Carpinteria, CA). Slides were then allowed to cool for 20 minutes in the same solution, rinsed in three changes of distilled water, and placed in Tris Buffered Saline with Tween 20 pH 7.4 (ScyTek Laboratories, Inc., West Logan, UT) for 5 minutes. The PolyVue HRP/DAB non-biotin polymer detection system (Diagnostic Biosystems, Pleasanton, CA) was used in the immunostaining protocols. Incubations occurred at room temperature unless otherwise specified, and for each step the sections were coated with 2-3 drops of reagent. Tris Buffered Saline with Tween 20 (pH 7.4) was used to rinse the sections between each step. Background Sniper solution (Biocare Medical, Concord, CA) was used to block non-specific staining for 20 minutes at room temperature. The mouse MSLN antibody (Rockland Immunochemicals, Inc. Antibodies & Assays, Gilbertsville, PA) was diluted (1:200) using the Renaissance Background Reducing Diluent solution (Biocare Medical). Slides were incubated with 100µl of the primary antibody solution for 1 hour at room temperature. Slides were then incubated in the universal secondary antibody provided with the kit. Afterwards, stable DAB Plus was applied for 1-2 minutes as a chromagen and slides were rinsed in distilled water, manually counterstained with Hematoxylin (Biocare Medical) for 10 seconds, and then rinsed in distilled water. Coverslips were then applied to each slide, using synthetic glass and permount mounting media. Negative and positive controls (negative; LNCaP, positive; OVCAR-3) were included in each immunostaining procedure.

## Data availability statement

Data generated during the study is maintained in electronic format on a database housed by the Center for Cell and Gene Therapy, Baylor College of Medicine. All data used in the analysis of the findings in the present study are included in the manuscript and the Supplementary Information. All other data (raw or analyzed) are available from the corresponding author (A.M.L.) on reasonable request.

## Results

### MSLN expression in cell lines and primary tumors

We evaluated the expression of MSLN across various cancers, both by flow cytometry and immunohistochemistry. As shown in **Supp. Figure 1A** MSLN is expressed at elevated levels in several cancer cell lines [mesothelioma (NCI-H2052 - 64%, NCI-H2452 - 94%), TNBC (HCC1806 - 87%), uterine (KLE - 95%), cervical (MS751 - 85%), colon (SK-CO-1 - 87%), ovarian (OVCAR3 - 76%), and pancreatic cancer cell lines (CAPAN-1 - 35% and CFPAC-1 - 31%)] and primary pancreatic tumor samples analyzed by immunohistochemistry (IHC), of which 7 of 9 were MSLN+ (**Supp. Figure 1B**). Negative controls include the cell lines HS578T, LNCaP and NCI-H28, representing TNBC, prostate cancer and mesothelioma, respectively (**Supp. Figure 1A**, right panel).

### Expansion, immunophenotype and specificity of MSLN-specific T cells

To next investigate the immunogenicity of MSLN to native T cells we exposed PBMCs from healthy donors (n = 29) with diverse HLA types (**Supp. Table 1**) to pepmix-loaded DCs, followed by expansion in T cell medium supplemented with Th1 polarizing, proproliferative, and prosurvival cytokines. After 23±2 days and 3 rounds of stimulation, 21 of the 29 lines exhibited anti-MSLN activity [defined as detection of >50 IFNγ spot forming cells (SFC)/2×10^5^ input cells], with a mean frequency of 1528±504 SFC/2×10^5^ input cells (median, 736; range 114 – 10,089 SFC in responders) with negligible non-specific activity (control: mean 18±2 SFC/2×10^5^ input cells, *P* < 0.005) (**Figure 1A**). In MSLN responders, a mean 32.3 ± 5.2-fold expansion in cell numbers (**Figure 1B**) was achieved yielding 276 ± 54×10^6^ total cells (**Supp. Figure 2**; **Supp. Table 2**). The expanded cells were comprised predominantly of CD3^+^ cells (97.6±1%) representing CD4^+^ (40.5±4.9%) and CD8^+^ (60.0±4.6%) subsets expressing both central [CD45RO^+^/CD62L^+^ (57.1±4.3%)] and effector memory markers [CD45RO^+^/CD62L^−^ (40.7±4.5%)], a composition poised for immediate effector function and long-term in vivo persistence. Furthermore, these cells were activated based on upregulation of CD25, CD28 and CD69 (64.7±2.3%, 46.7±4.4% and 40.5±2.3%, respectively), with minimal co-expression of exhaustion markers (CD3^+^/PD1^+^/TIM3^+^/LAG3^+^: 1.8±0.3%, **Figure 1C**).

**Figure 1.**
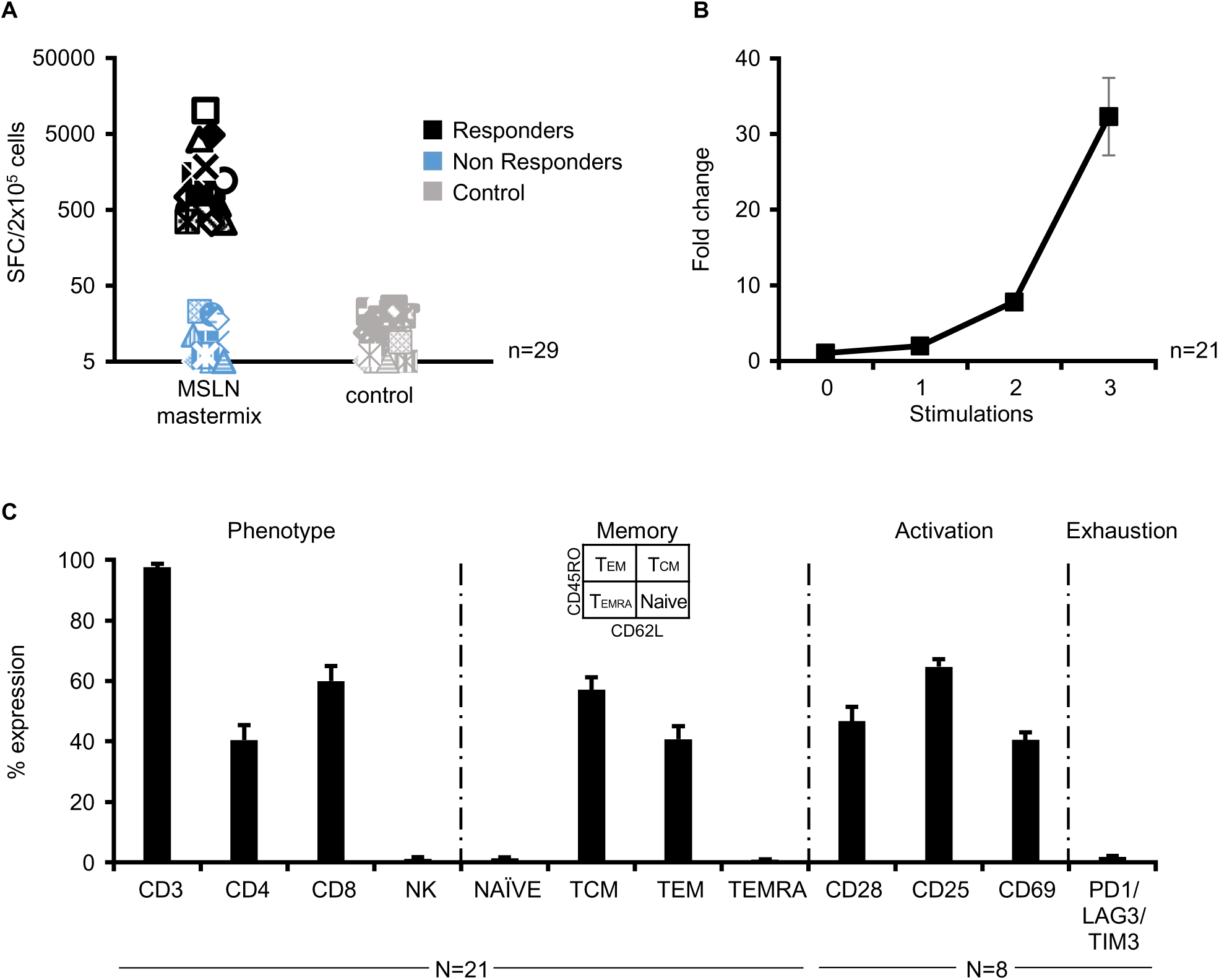
MSLN-specific T cell specificity, expansion and immunophenotypic profile. (A) Specificity of generated cell lines (n = 29) evaluated by IFNγ ELISpot. 21 of the 29 lines exhibited anti-MSLN activity (defined as detection of >50 SFC/2×10^5^ input cells). (B) Mean (± SEM) fold expansion of MSLN-specific T cells in the responder cohort (n=21). (C) Phenotype, memory/activation profile and exhaustion markers of MSLN-specific T cells in responders, reported as % expression (mean±SEM, n=8-21). MSLN, mesothelin; ELISpot, enzyme-linked immunospot; IFNγ, interferon gamma; SFC, spot-forming cells.

### Ex vivo-expanded MSLN-specific T cells are polyfunctional

We next examined whether both CD4^+^ and CD8^+^ T cells contained MSLN-reactive cells by performing ICS for IFNγ and TNFα. MSLN-directed activity was predominantly detected in the CD8^+^ compartment with weaker CD4 reactivity (**Figure 2A**, left panel - representative results from 2 donors; **Figure 2A**, right panel - summary data, 17±4.1% CD8^+^/IFNγ-producing cells; 3.5±2% CD4^+^/IFNγ-producing cells, n = 21). The in vivo efficacy of infused T cells has been closely linked to their capacity to produce multiple effector molecules (i.e., polyfunctionality), and we were encouraged to find that the majority of reactive cells produced both IFNγ and TNFα, as assessed by ICS (**Figure 2B**, left panel - representative results from 2 donors; **Figure 2B**, right panel - summary data, 14.8±3.7% CD8^+^/IFNγ/TNFα-producing cells; 1.8±1.2% CD4^+^/IFNγ/TNFα-producing cells, n = 21). When we further explored polyfunctionality using the FluoroSpot assay we found that following MSLN exposure reactive T cells secreted not only IFNγ (810±261 SFC/2×10^5^ cells) and TNFα (780±354 SFC/2×10^5^ cells) but also granzyme B (GrB; 444±158 SFC/2×10^5^ cells) (**Figure 2C**, top). Furthermore, of all IFNγ-producing cells, 52% were polyfunctional, additionally producing either GrB (8%), TNFα (36%) or both GrB and TNFα (8%) (n=6). These results were corroborated when focusing on TNFα or GrB (61% and 57% polyfunctional, respectively; **Figure 2C**, bottom).

**Figure 2.**
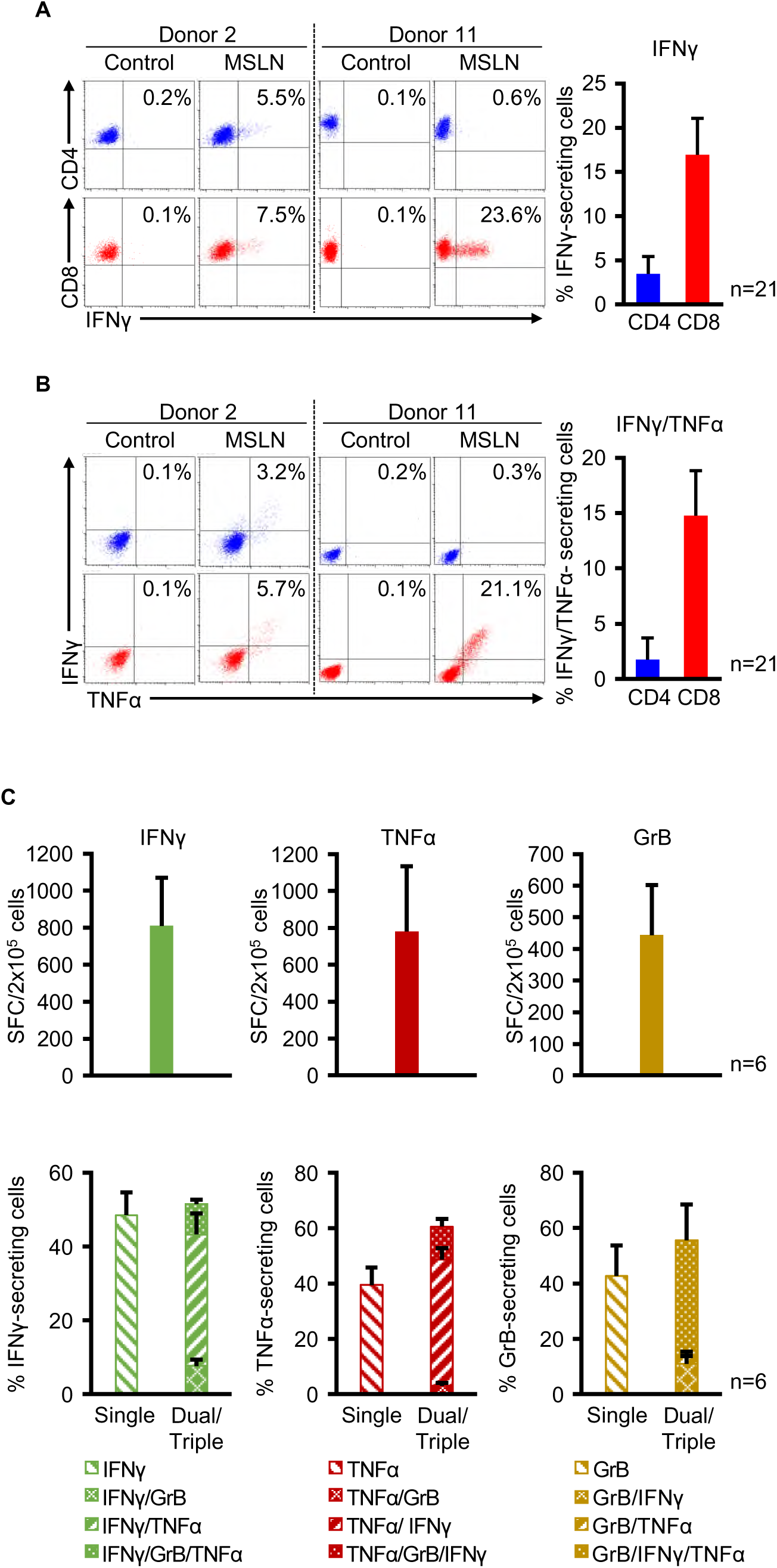
Effector functions of MSLN-specific T cells. (A) Representative donor data (left panel) and summary results (mean±SEM, n=21; right panel) showing CD8⁺/IFNγ⁺ and CD4⁺/IFNγ⁺ T cells, as assessed by ICS. (B) Representative donor data (left panel) and summary results (mean±SEM; n=21; right panel) showing dual (IFNγ/TNFα)-producing CD4⁺ and CD8⁺ MSLN-specific T cells, as assessed by ICS. (C) Polyfunctionality of MSLN-specific T cells as measured by FluoroSpot (IFNγ/Granzyme B/TNFα) (n=6). Top panel shows the total frequency (SFC/2×10^5^ cells) of specific T cells for each of the individual effector molecules, while the bottom panel shows the proportion of cells that are single, dual or triple analyte-producing. TNFα, tumor necrosis factor alpha; ICS, intracellular cytokine staining; GrB, Granzyme B.

Though FluoroSpot (**Figure 2C**) and ELISpot (**Supp. Figure 3**) GrB results were indicative of cytolytic potential, we formally examined lytic activity by incubating MSLN-reactive T cells with antigen-loaded autologous (PHA blasts) or control non-antigen-expressing targets. T cells were able to specifically lyse autologous MSLN-loaded targets (**Figure 3A** - representative results from 2 donors; **Figure 3B** - summary data, median, 55% specific lysis; range 26-90%, 40:1 E:T, n=17, *P* <0.0005), and killing was diminished in the presence of MHC class I but not MHC class II blocking antibodies. Furthermore, reactive cells were also able to recognize and kill partially HLA-matched cell lines that endogenously expressed MSLN, as shown in **Figure 3C**.

**Figure 3.**
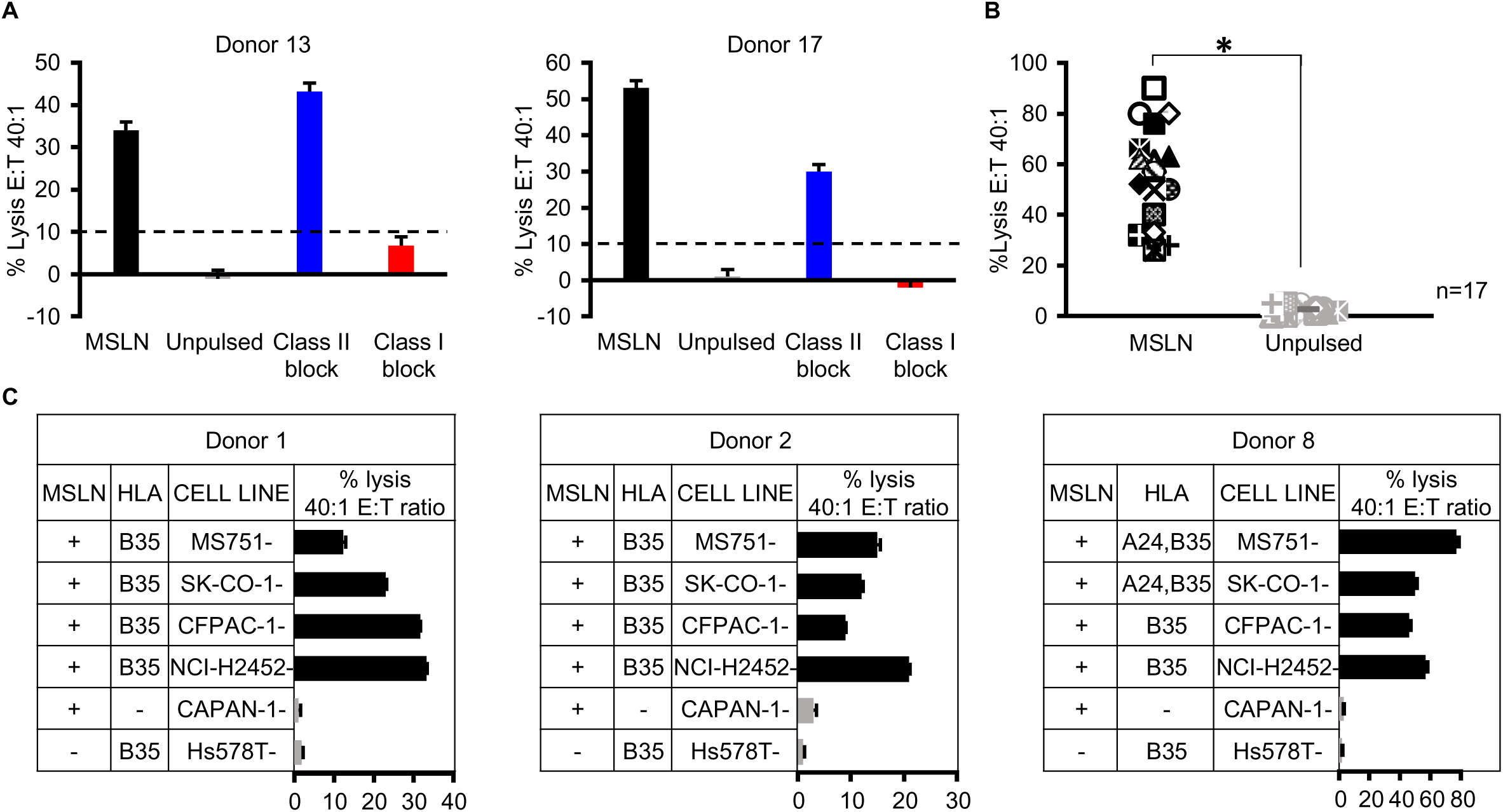
Cytolytic capacity of MSLN-specific T cells in 2D cultures. (A) Representative donor data showing lysis of MSLN-loaded autologous PHA blasts in the absence or presence of MHC class I or class II blocking antibodies (triplicates, mean ± SD). (B) Summary cytotoxicity data (n = 17). (C) Killing of cell lines endogenously expressing MSLN by partially HLA-matched MSLN-specific T cells from three independent donors (triplicates, mean ± SD). Cytotoxicity in all panels was assessed by ⁵¹Cr release assay following a 5–8 h co-culture at a 40:1 effector-to-target (E:T) ratio. PHA, phytohemagglutinin; MHC, major histocompatibility complex.

3D culture systems more closely replicate in vivo characteristics with tumor spheroids providing a model that better mimics the solid tumor architecture and physiology^33, 34, 35^. Hence, we next explored the benefits of our strategy with MSLN+ pancreatic cancer and cervical cancer-derived 3D tumor spheroids. We first established spheroids using GFP/FFLuc-labeled tumor cells (**Figure 4A**). To assess anti-tumor activity we co-cultured tumor spheroids alone (Tumor), or with partially HLA-matched MSLN-specific or control T cells at the indicated E:T (based on initial number of tumor cells used to generate spheroids). Bioluminescence imaging of the spheroids demonstrated a decline in tumor burden in both MSLN T cell-treated conditions (**Figure 4B-C**). Taken together these results demonstrate that MSLN-specific T cells can recognize and potently kill MSLN-expressing targets in 2D and 3D culture systems.

**Figure 4.**
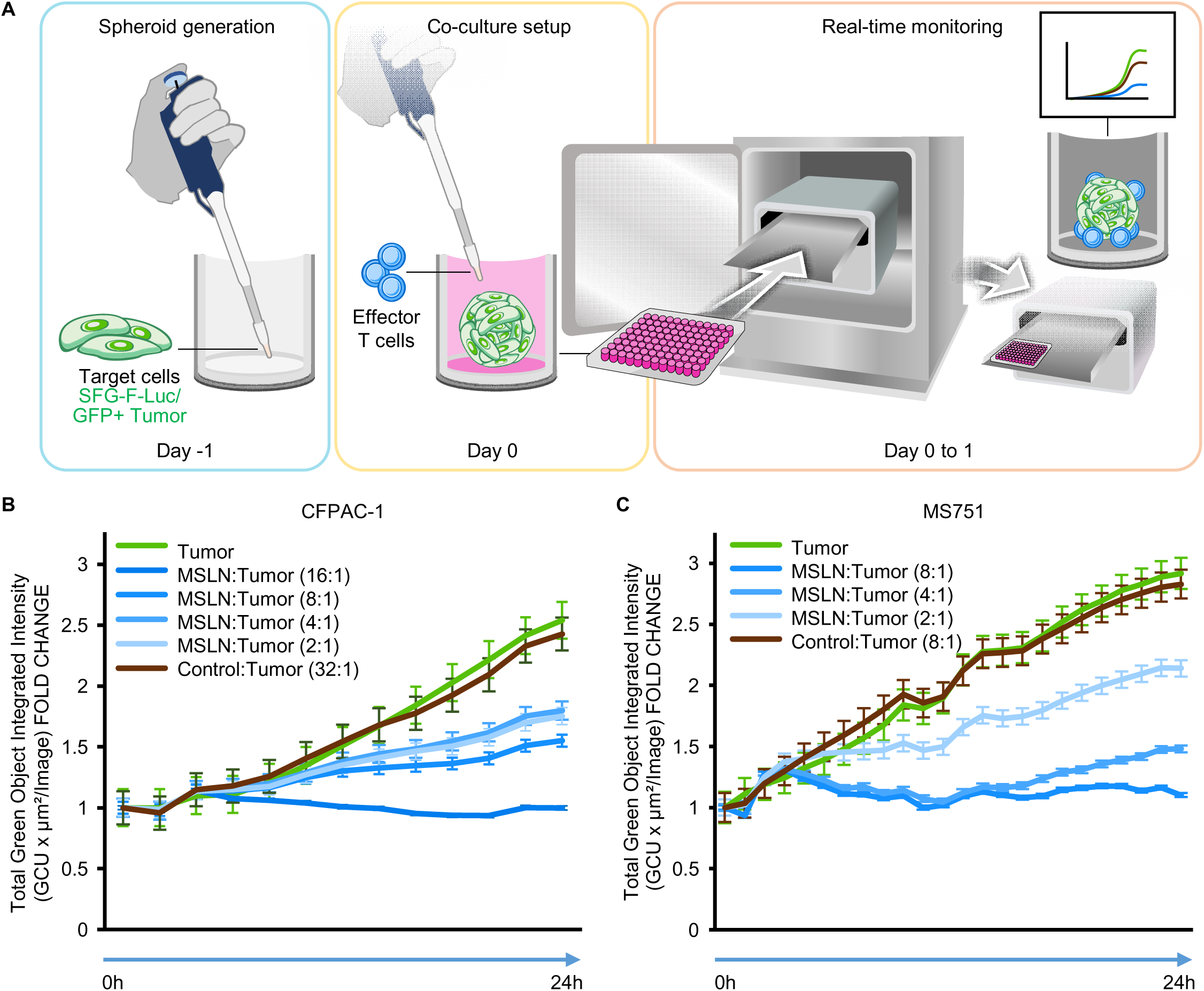
Cytolytic capacity of MSLN-specific T cells in 3D tumor models. (A) Illustration of the experimental procedure for the 3D co-culture assay using GFP/FFluc+ tumor spheroids and effector T cells. (B-C) Tumor spheroid bioluminescence in the absence of T cells (Tumor), and in the presence of MSLN or Control T cells at the indicated E:T ratios. Tumor spheroids were derived from the CFPAC-1 (B) and MS751 (C) cell lines.

### Magnitude and specificity of MSLN reactivity within ex vivo-expanded T cell lines

To better understand the breadth of T cell epitope reactivity within our polyclonal T cell lines, we utilized an overlapping peptide library comprised of 153 15 mer peptides (**Supp. Table 3**) spanning the entire antigen sequence. These peptides were divided into 25 minipools, each containing 12 or 13 peptides and organized such that each individual peptide was present in 2 minipools. Then reactive T cell lines were rescreened using these minipools. An example of the screening process is shown in **Figure 5A**, in which a T cell line generated from donor 15 reacted against minipools 1, 6, 15, and 25, revealing 4 potential stimulatory peptides - #25, #30, #145 and #150 (**Figure 5B**). Subsequent screening with each of these individual peptides by ICS for IFNγ revealed the two epitope-containing peptides as #30 and #145 (**Figure 5C**). **Supp. Figure 4 (A-C)** shows similar data for donor 11 whose minipool reactivity mapped to pools 6, 8, 9, 15 and 23, which revealed epitope-specific responses in peptides # 30, #128 and #129.

**Figure 5.**
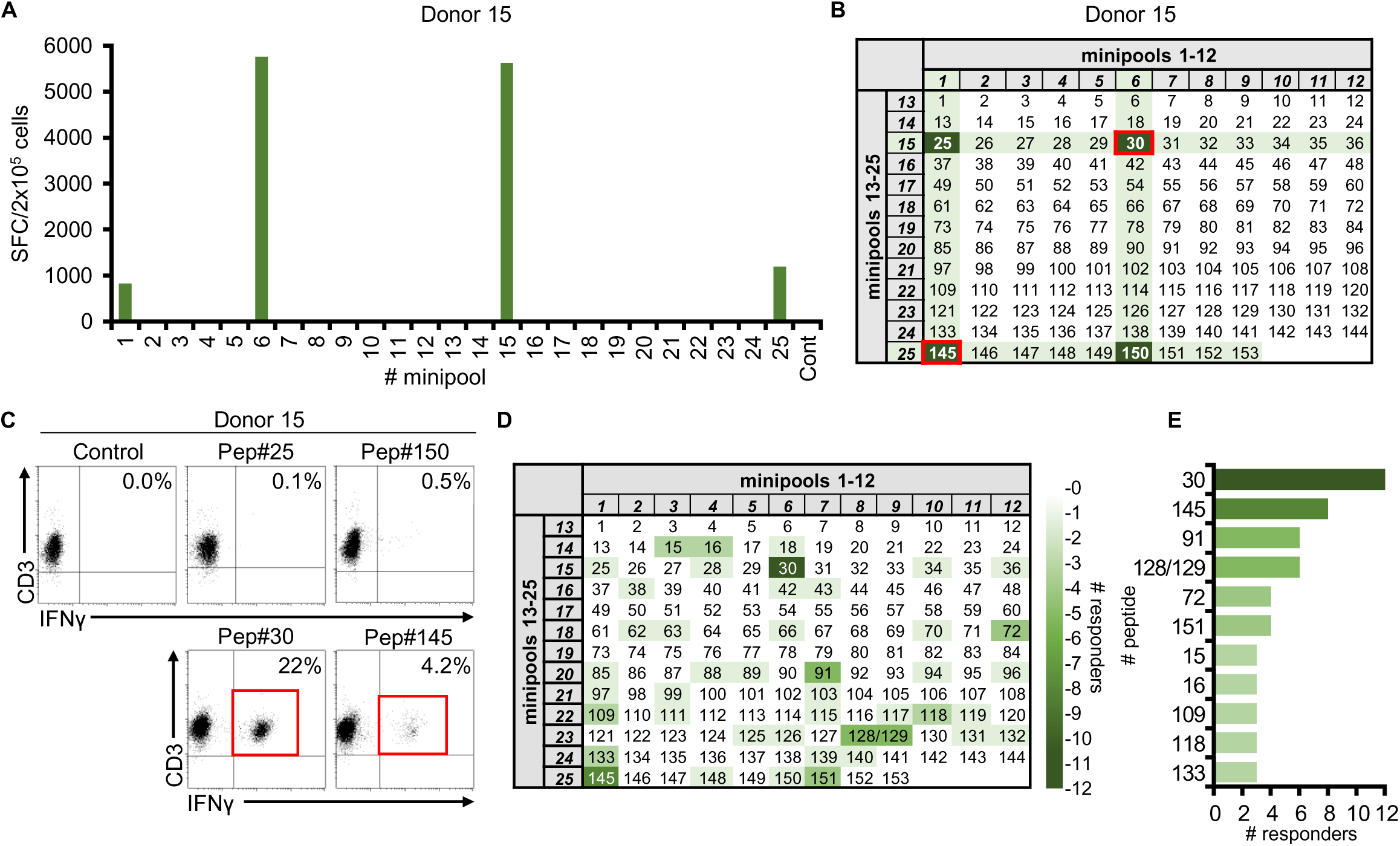
MSLN epitope identification. (A) To identify immunogenic peptide epitopes T cell lines were screened against individual MSLN minipools by IFNγ ELISpot. Panel A shows results for donor 15 (results reported as SFC/2×10^5^), with the potential stimulatory peptides shown in panel B. (C) To identify the immunogenic epitopes the MSLN T cell line was exposed to each of the peptides identified in panel B – screening was performed by ICS for IFNγ and results reported as % of CD3+/IFNγ+ cells. (D) Summary of identified immunogenic MSLN peptides across all 21 cell lines and color coded to identify the number of responders, with immunodominant peptides (found to elicit responses in ≥3 donors) shown in panel E.

Figure 5D summarizes the screening results for all 21 responding donors. Overall, 44 of the 153 peptides (29%) elicited a response in at least 1 donor and 11 peptides were recognized by ≥ 3 individuals (Figure 5E). Of these, peptide #30 was the most immunogenic, with reactivity identified in 12 individuals, followed in descending order by #145 (n=8), #91 and #128/129 (n=6), # 72 and 151 (n=4) and #s 15, 16, 109, 118 and 133 recognized by n=3. Of the most frequently recognized peptides, reactivity was primarily CD8-mediated, with the exception of peptide 72 against which responses were detected in both the CD4^+^ and CD8^+^ T cell compartment [**Supp. Figure 5 (A-D)**]. Of note, reactivity against peptides #109 and 133 reflected reactivity toward the previously described A24-restricted FYPGYLCSL^36, 37^ and A2-restricted VLPLTVAEV^37^ epitopes, respectively (**Table 1**).

**Table 1.**
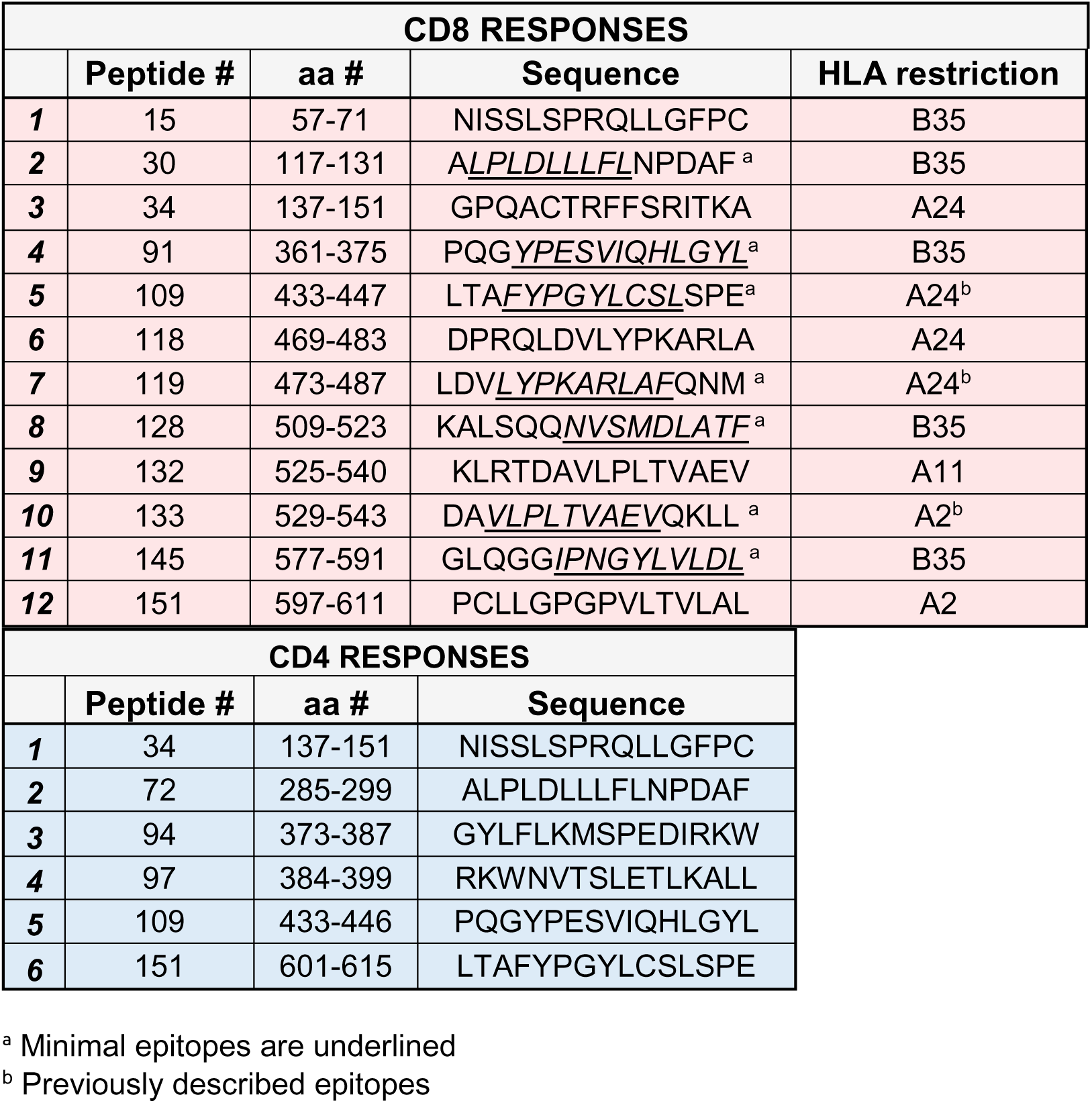
Immunogenic peptides identified within the CD4+ and the CD8+ compartment.

### Identifying the HLA-restricting alleles presenting novel T cell epitopes

We next sought to identify the HLA restricting alleles for the novel immunogenic peptides revealed by our screening studies, with specific focus on those that were immunodominant. Figure 6 **(A, B**, left panels), shows an example of the mapping studies performed on donor 15’s line (HLA-A: 24, 26; HLA-B: 35, 40), whose response to peptides #30 and #145 was CD8^+^ mediated, as revealed by ICS. To next identify the HLA-restricting allele peptide-pulsed autologous, and partially-HLA matched PHA blasts were used as targets in a cytotoxicity assay and only B35-matched peptide-pulsed targets were killed in each case [Figure 6 **(A, B**), right panels]. Similarly, peptides #91 and #128/129 were found to be presented through HLA-B35 (Figure 6C, **D**), while peptides #132, #151, #34 and #118 mapped to HLA-A11, HLA-A2, HLA-A24 and HLA-A24, respectively [(**Supp. Figure 6 (A-D**)]. Finally, to identify the minimal epitopes recognized by peptides #30, 145, 91 and 128/129, we used a panel of 9-14 mers, which revealed the minimal epitopes as 9mer - LPLDLLLFL (**Supp. Figure 7A**), 9mer – IPNGYLVLD (**Supp. Figure 7B**), 12mer – YPESVIQHLGYL (**Supp. Figure 7C**) and 9mer – NVSMDLATF (**Supp. Figure 7D**) respectively. **Table 1** summarizes all stimulatory peptides identified, with mapped minimal epitopes underlined as well as HLA restrictions that have been mapped to date^36, 37^.

**Figure 6.**
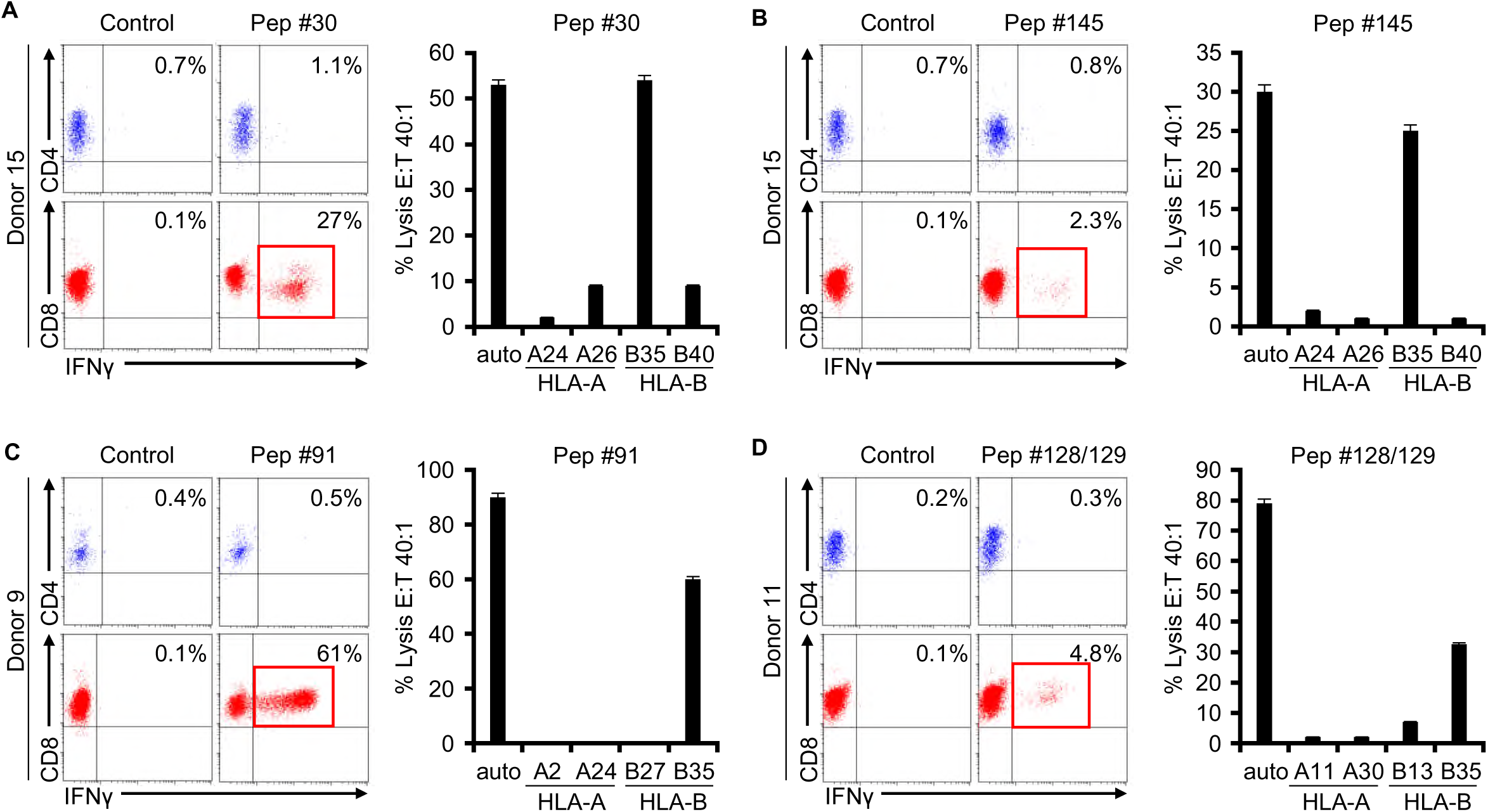
Identification of HLA-restricting alleles for the immunodominant MSLN peptides. (A) For peptide 30: ICS (IFNγ production) analysis to determine whether reactive T cells were detected in the CD4+ or CD8+ T cell compartment (left panel). ⁵¹Cr release assay (5-8 hr co-culture) using peptide-pulsed autologous and partially HLA-matched PHA blasts as targets (right panel). Results show % specific lysis (mean±SD), E:T 40:1. Panels B-D show similar data for peptides 145, 91, and 128/129, respectively. HLA, Human Leukocyte Antigen.

## Discussion

Targeting solid tumors remains a hurdle for immunotherapeutic applications due to a range of issues, one of which is the dearth of suitable target antigens exclusively expressed/overexpressed on tumor cells. In the current study we explored the immunotherapeutic potential of MSLN – an antigen that is differentially expressed in several cancer types, many with poor prognosis. Both its overexpression and its contribution to maintaining the malignant phenotype make it an attractive target, and here we evaluated the potential of targeting MSLN-expressing tumor cells using T cells with native TCR specificity. We demonstrate that MSLN-specific T cells can be readily expanded from donors of diverse HLA backgrounds, with dominant activity detected in the CD8^+^ T cell subset. Reactive cells are Th1 polarized, polyfunctional, and capable of mediating potent cytotoxic effects, resulting in pronounced anti-tumor effects against mesothelioma, cervical, colon and pancreatic cancer cells in 2D and 3D tumor models. Taken together, these data support the clinical translation of these cells, either as a single agent or combined with additional specificities to enhance activity against heterogeneous tumors.

MSLN is a GPI-anchored cell surface protein that is overexpressed in many different solid tumor types, including mesothelioma, ovarian and pancreatic cancer, gastric adenocarcinoma and TNBC^7, 10, 11, 12, 13, 21, 38^. In general GPI-linked proteins play a role in cell signaling and adhesion and although the physiologic function of MSLN is unclear, in the malignant setting there is evidence that it plays a role in promoting tumor cell proliferation and dissemination^9, 14, 16, 18, 19, 20^. In an effort to therapeutically exploit MSLN a number of targeted modalities have been developed including vaccines^39^, as well as agents targeted to surface-expressed MSLN such as monoclonal antibodies, immunotoxins, antibody-drug conjugates^5, 22, 23, 24, 25^ and CAR T cells^26, 27, 28, 29, 30, 31, 32^. However, cell surface cleavage and the potential that soluble MSLN (present at elevated levels in the sera of patients with solid tumors) may interfere with recognition is a consideration^40^. In contrast, adoptively transferred T cells with native TCR specificity recognize target antigen that has been processed and presented on the cell surface in the context of HLA molecules and as such are impervious to both the presence of soluble antigen as well as cleavage events.

Jaffee and colleagues^41^ were the first to identify that MSLN was immunogenic to endogenous T cells in correlative studies performed on patients in receipt of GM-CSF-secreting pancreatic cancer cell lines. Indeed, in patients who developed post-vaccination delayed-type hypersensitivity (DTH) responses they discovered amplified levels of T cells directed to epitopes presented in the context of HLA-A2, A3, and A24^37^. Furthermore, in subsequent Phase I and II studies of patients with PDAC, prolonged disease-free survival was shown to correlate with the induction and maintenance of elevated levels of MSLN-directed T cells in HLA-A1, HLA-A2 and HLA-A3 positive patients^42, 43, 44, 45^. In the current study we sought to expand the immune library of MSLN-targeted epitopes using an unbiased approach in which we used overlapping peptides (153 15mers, each overlapping by 11 amino acids) to interrogate both CD4^+^ and CD8^+^ T cell epitopes in individuals of diverse HLA backgrounds. Overall we found immune responses mapping to almost one third of the antigen (44/153 peptides elicited a response in at least 1 donor) and in addition to detecting several previously documented CD8 responses^36,37^, we mapped numerous novel CD8^+^ T cell responses restricted to peptides presented in the context of HLA-B35, A2, A11 and A24. Though CD4^+^ responses were less frequently detected, we were able to identify the presence of overlapping CD4^+^ and CD8^+^ T cell epitopes within peptide #72 (aa ERTILRPRFRREVEK), which was recognized by 4 of our screened donors.

Our group has clinically targeted a range of cancer testis/shared/differentiation antigens including PRAME, SSX2, MAGEA4, SURVIVIN, NY-ESO-1, and WT-1 in diseases including leukemia^46, 47^, lymphoma^48^, multiple myeloma^49^, breast^50^ and pancreatic cancer (in press) using ex vivo expanded autologous or allogeneic T cells without severe infusion-related toxicities and with documented long-term clinical effects. Our current work supports extending our target range to include MSLN for a broad range of solid tumors. MSLN is immunogenic, and it is feasible to generate autologous MSLN-specific T cells that recognize antigen-loaded and endogenously expressing target cells. This, in addition to previous reports documenting an association between endogenous MSLN-specific T cells in vaccine recipients and prolonged clinical benefit. These findings support targeting MSLN (alone or in combination with other tumor-expressed antigens) in future clinical trials of adoptively transferred T cells for solid tumors.

## Acknowledgments

The authors would like to acknowledge Walter M. Mejia for help with figure generation and manuscript formatting. This work was supported by a Translational Research Award from the V Foundation (PI: A.M.L.) and the Dan L. Duncan Comprehensive Cancer Center for application of the shared resources from a support grant from the National Institutes of Health, National Cancer Institute (P30CA125123).

## Notes

### Competing Interest Statement

The authors have declared no competing interest.

