## Supplementary data for "Targeting Mesothelin via the native T cell receptor"

|  | **# DONOR** | **HLA-A** | **HLA-B** | **HLA- DR** | **HLA- DQ** |
| --- | --- | --- | --- | --- | --- |
| **Responders** | 1 | 02, 11 | 35, 38 | 10, 13 | 01, 01 |
|  | 2 | 02, 30 | 35, 13 | 07, 08 | 02, 06 |
|  | 3 | 24, 11 | 35, 40 | 04, 14 | 04, 05 |
|  | 4 | 11, 23 | 35, 57 | 01, 07 | N/D |
|  | 5 | 02, 24 | 35, 13 | 12, 15 | 03, 06 |
|  | 6 | 02, 24 | 40, 44 | 11, 14 | 03, 05 |
|  | 7 | 02,25 | 15,51 | 04, 08 | 03, 04 |
|  | 8 | 24, 24 | 35, 35 | 07, 08 | 02, 04 |
|  | 9 | 02, 24 | 35, 27 | 01, 11 | 03, 05 |
|  | 10 | 01, 68 | 35, 08 | 03, 11 | 02, 03 |
|  | 11 | 11, 30 | 35, 13 | 01, 07 | 02, 05 |
|  | 12 | 03, 24 | 35, 08 | 03, 04 | 02, 03 |
|  | 13 | 11,24 | 35, 55 | 08, 09 | N/D |
|  | 14 | 02, 31 | 07, 40 | 04, 15 | 03, 06 |
|  | 15 | 24, 26 | 35, 40 | 08, 12 | 03, 03 |
|  | 16 | 11, 33 | 35, 39 | 04, 08 | 03, 06 |
|  | 17 | 03, 01 | 35, 08 | 03, 11 | 02, 03 |
|  | 18 | 02, 31 | 35, 40 | 04, 14 | 04, 05 |
|  | 19 | 11, 68 | 35, 53 | N/D | N/D |
|  | 20 | 02, 03 | 35, 46 | 04, 09 | 01, 03 |
|  | 21 | 02, 03 | 13, 35 | 04, 11 | 05, 08 |
| **Non responders** | 22 | 02, 24 | 15, 35 | 09, 12 | 03, 03 |
|  | 23 | 02, 33 | 40, 44 | 09, 13 | 03, 06 |
|  | 24 | 11 | 46 | 09 | 03 |
|  | 25 | 02, 62 | 15 | 04 | 03,03 |
|  | 26 | 01, 11 | 08, 49 | 08, 13 | 04, 06 |
|  | 27 | 01, 02 | 51, 57 | 04, 13 | 03, 06 |
|  | 28 | 01, 26 | 40 | 07, 15 | 02, 06 |
|  | 29 | 02, 68 | 15, 52 | 04, 15 | 03, 06 |

| **Expansion characteristics** | | | |
| --- | --- | --- | --- |
| **Donor** | **#cells (x10^6^)** | | **Fold change** |
|  | **Day0** | **Day23** |  |
| *1* | 6 | 171 | 29 |
| *2* | 4 | 202 | 50 |
| *3* | 4 | 53 | 13 |
| *4* | 6 | 45 | 7 |
| *5* | 3 | 132 | 44 |
| *6* | 4 | 56 | 14 |
| *7* | 6 | 46 | 8 |
| *8* | 6 | 63 | 11 |
| *9* | 10 | 496 | 50 |
| *10* | 10 | 211 | 21 |
| *11* | 10 | 211 | 21 |
| *12* | 10 | 201 | 20 |
| *13* | 10 | 253 | 25 |
| *14* | 10 | 173 | 17 |
| *15* | 10 | 253 | 25 |
| *16* | 10 | 291 | 29 |
| *17* | 10 | 409 | 41 |
| *18* | 10 | 210 | 21 |
| *19* | 10 | 887 | 89 |
| *20* | 10 | 530 | 53 |
| *21* | 10 | 908 | 91 |

| **#**  **pep** | **Sequence** | **#**  **pep** | **Sequence** | **#**  **pep** | **Sequence** | **#**  **pep** | **Sequence** |
| --- | --- | --- | --- | --- | --- | --- | --- |
| *1* | MALPTARPLLGSCGT | *40* | PERQRLLPAALACWG | *79* | ESLIFYKKWELEACV | *118* | DPRQLDVLYPKARLA |
| *2* | TARPLLGSCGTPALG | *41* | RLLPAALACWGVRGS | *80* | FYKKWELEACVDAAL | *119* | LDVLYPKARLAFQNM |
| *3* | LLGSCGTPALGSLLF | *42* | AALACWGVRGSLLSE | *81* | WELEACVDAALLATQ | *120* | YPKARLAFQNMNGSE |
| *4* | CGTPALGSLLFLLFS | *43* | CWGVRGSLLSEADVR | *82* | ACVDAALLATQMDRV | *121* | RLAFQNMNGSEYFVK |
| *5* | ALGSLLFLLFSLGWV | *44* | RGSLLSEADVRALGG | *83* | AALLATQMDRVNAIP | *122* | QNMNGSEYFVKIQSF |
| *6* | LLFLLFSLGWVQPSR | *45* | LSEADVRALGGLACD | *84* | ATQMDRVNAIPFTYE | *123* | GSEYFVKIQSFLGGA |
| *7* | LFSLGWVQPSRTLAG | *46* | DVRALGGLACDLPGR | *85* | DRVNAIPFTYEQLDV | *124* | FVKIQSFLGGAPTED |
| *8* | GWVQPSRTLAGETGQ | *47* | LGGLACDLPGRFVAE | *86* | AIPFTYEQLDVLKHK | *125* | QSFLGGAPTEDLKAL |
| *9* | PSRTLAGETGQEAAP | *48* | ACDLPGRFVAESAEV | *87* | TYEQLDVLKHKLDEL | *126* | GGAPTEDLKALSQQN |
| *10* | LAGETGQEAAPLDGV | *49* | PGRFVAESAEVLLPR | *88* | LDVLKHKLDELYPQG | *127* | TEDLKALSQQNVSMD |
| *11* | TGQEAAPLDGVLANP | *50* | VAESAEVLLPRLVSC | *89* | KHKLDELYPQGYPES | *128* | KALSQQNVSMDLATF |
| *12* | AAPLDGVLANPPNIS | *51* | AEVLLPRLVSCPGPL | *90* | DELYPQGYPESVIQH | *129* | QQNVSMDLATFMKLR |
| *13* | DGVLANPPNISSLSP | *52* | LPRLVSCPGPLDQDQ | *91* | PQGYPESVIQHLGYL | *130* | SMDLATFMKLRTDAV |
| *14* | ANPPNISSLSPRQLL | *53* | VSCPGPLDQDQQEAA | *92* | PESVIQHLGYLFLKM | *131* | ATFMKLRTDAVLPLT |
| *15* | NISSLSPRQLLGFPC | *54* | GPLDQDQQEAARAAL | *93* | IQHLGYLFLKMSPED | *132* | KLRTDAVLPLTVAEV |
| *16* | LSPRQLLGFPCAEVS | *55* | QDQQEAARAALQGGG | *94* | GYLFLKMSPEDIRKW | *133* | DAVLPLTVAEVQKLL |
| *17* | QLLGFPCAEVSGLST | *56* | EAARAALQGGGPPYG | *95* | LKMSPEDIRKWNVTS | *134* | PLTVAEVQKLLGPHV |
| *18* | FPCAEVSGLSTERVR | *57* | AALQGGGPPYGPPST | *96* | PEDIRKWNVTSLETL | *135* | AEVQKLLGPHVEGLK |
| *19* | EVSGLSTERVRELAV | *58* | GGGPPYGPPSTWSVS | *97* | RKWNVTSLETLKALL | *136* | KLLGPHVEGLKAEER |
| *20* | LSTERVRELAVALAQ | *59* | PYGPPSTWSVSTMDA | *98* | VTSLETLKALLEVNK | *137* | PHVEGLKAEERHRPV |
| *21* | RVRELAVALAQKNVK | *60* | PSTWSVSTMDALRGL | *99* | ETLKALLEVNKGHEM | *138* | GLKAEERHRPVRDWI |
| *22* | LAVALAQKNVKLSTE | *61* | SVSTMDALRGLLPVL | *100* | ALLEVNKGHEMSPQV | *139* | EERHRPVRDWILRQR |
| *23* | LAQKNVKLSTEQLRC | *62* | MDALRGLLPVLGQPI | *101* | VNKGHEMSPQVATLI | *140* | RPVRDWILRQRQDDL |
| *24* | NVKLSTEQLRCLAHR | *63* | RGLLPVLGQPIIRSI | *102* | HEMSPQVATLIDRFV | *141* | DWILRQRQDDLDTLG |
| *25* | STEQLRCLAHRLSEP | *64* | PVLGQPIIRSIPQGI | *103* | PQVATLIDRFVKGRG | *142* | RQRQDDLDTLGLGLQ |
| *26* | LRCLAHRLSEPPEDL | *65* | QPIIRSIPQGIVAAW | *104* | TLIDRFVKGRGQLDK | *143* | DDLDTLGLGLQGGIP |
| *27* | AHRLSEPPEDLDALP | *66* | RSIPQGIVAAWRQRS | *105* | RFVKGRGQLDKDTLD | *144* | TLGLGLQGGIPNGYL |
| *28* | SEPPEDLDALPLDLL | *67* | QGIVAAWRQRSSRDP | *106* | GRGQLDKDTLDTLTA | *145* | GLQGGIPNGYLVLDL |
| *29* | EDLDALPLDLLLFLN | *68* | AAWRQRSSRDPSWRQ | *107* | LDKDTLDTLTAFYPG | *146* | GIPNGYLVLDLSMQE |
| *30* | ALPLDLLLFLNPDAF | *69* | QRSSRDPSWRQPERT | *108* | TLDTLTAFYPGYLCS | *147* | GYLVLDLSMQEALSG |

| *31* | DLLLFLNPDAFSGPQ | *70* | RDPSWRQPERTILRP | *109* | LTAFYPGYLCSLSPE | *148* | LDLSMQEALSGTPCL |
| --- | --- | --- | --- | --- | --- | --- | --- |
| *32* | FLNPDAFSGPQACTR | *71* | WRQPERTILRPRFRR | *110* | YPGYLCSLSPEELSS | *149* | MQEALSGTPCLLGPG |
| *33* | DAFSGPQACTRFFSR | *72* | ERTILRPRFRREVEK | *111* | LCSLSPEELSSVPPS | *150* | LSGTPCLLGPGPVLT |
| *34* | GPQACTRFFSRITKA | *73* | LRPRFRREVEKTACP | *112* | SPEELSSVPPSSIWA | *151* | PCLLGPGPVLTVLAL |
| *35* | CTRFFSRITKANVDL | *74* | FRREVEKTACPSGKK | *113* | LSSVPPSSIWAVRPQ | *152* | GPGPVLTVLALLLAS |
| *36* | FSRITKANVDLLPRG | *75* | VEKTACPSGKKAREI | *114* | PPSSIWAVRPQDLDT | *153* | VLTVLALLLASTLA |
| *37* | TKANVDLLPRGAPER | *76* | ACPSGKKAREIDESL | *115* | IWAVRPQDLDTCDPR |  | |
| *38* | VDLLPRGAPERQRLL | *77* | GKKAREIDESLIFYK | *116* | RPQDLDTCDPRQLDV |  |  |
| *39* | PRGAPERQRLLPAAL | *78* | REIDESLIFYKKWEL | *117* | LDTCDPRQLDVLYPK |  |  |

**A**	MSLN+

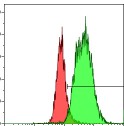

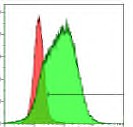

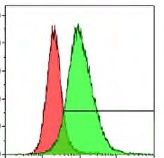

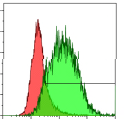

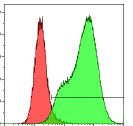

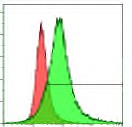

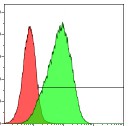

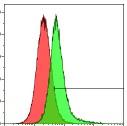

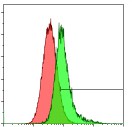

**31%**

**87%**

**87%**

**CFPAC-1**

**SK-CO-1**

**HCC1806**

**35%**

**85%**

**94%**

**CAPAN-1**

**MS751**

**NCI-H2452**

**76%**

**95%**

**64%**

**OVCAR-3**

**KLE**

**NCI-H2052**

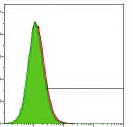

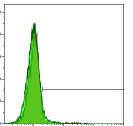

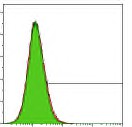

**HS578T**

**0.4%**

**LNCaP**

**0.1%**

**NCI-H28**

**0.0%**

MPM

MPM

Cervical cancer

Uterine cancer

PDAC

Ovarian cancer

Prostate cancer

TNBC

MSLN-

**B**	MSLN-

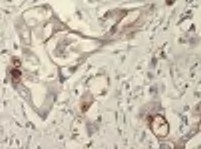

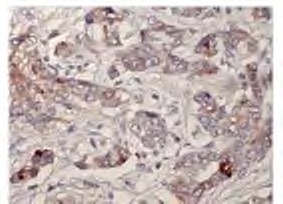

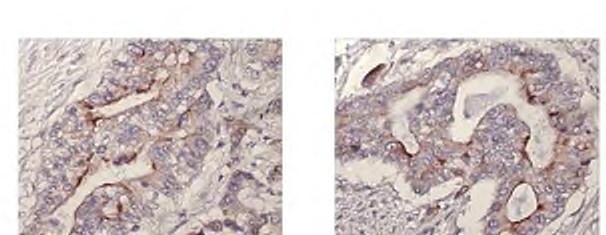

MSLN+

TNBC

PDAC

Colon cancer

PDAC

MPM

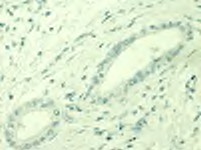

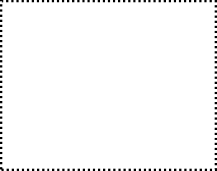

**Supp. figure 1. MSLN expression in cell lines and primary tumors.** (A) Flow cytometric evaluation of MSLN expression in several cancer cell lines [mesothelioma (NCI-H2052, NCI-H2452), TNBC (HCC1806), uterine (KLE), cervical (MS751), colon (SK-CO-1), ovarian (OVCAR3), and pancreatic cancer (CAPAN-1 and CFPAC-1)] ((left panel). Negative controls (HS578T, LNCaP and NCI-H28, representing TNBC, prostate cancer and mesothelioma, respectively) are also shown (right panel). ((B) Assessment of MSLN expression in primary pancreatic tumor samples by immunohistochemistry (IHC). Representative examples of MSLN+ (left panel) and MSLN- (right panel) staining are shown.

350

300

250

\# Cells (x10^6^)

200

150

100

50

0

0	1	2	3

Stimulations

n=21

**Supp. figure 2. Ex vivo expansion of MSLN-specific T cells.** Cell numbers (mean ± SEM) achieved in the responder cohort (n=21) following each round of stimulation.

800

700

600

SFC/2x10^5^ cells

500

400

300

200

100

0

Granzyme B ELISpot

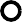

MSLN	control	n=6

**Supp. figure 3. Effector functions of MSLN-specific T cells.** Production of Granzyme B by MSLN-STs following direct antigenic stimulation as assessed by ELISpot; results are reported as SFC/2x10^5^ input cells. Individual values are shown (n=6).

**A**	1500

SFC /2x10^5^ cells

1000

**Donor 11**

**C	IFNγ production**

Control

500

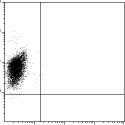

0.1%

0

1

2

3

4

5

6

7

8

9

10

11

12

13

14

15

16

17

18

19

20

21

22

23

24

25

Cont

**# minipool**

**B**

|  | | **minipools 1-12** | | | | | | | | | | | |
| --- | --- | --- | --- | --- | --- | --- | --- | --- | --- | --- | --- | --- | --- |
|  |  | ***1*** | ***2*** | ***3*** | ***4*** | ***5*** | ***6*** | ***7*** | ***8*** | ***9*** | ***10*** | ***11*** | ***12*** |
| **minipools 13-25** | ***13*** | 1 | 2 | 3 | 4 | 5 | 6 | 7 | 8 | 9 | 10 | 11 | 12 |
|  | ***14*** | 13 | 14 | 15 | 16 | 17 | 18 | 19 | 20 | 21 | 22 | 23 | 24 |
|  | ***15*** | 25 | 26 | 27 | 28 | 29 | **30** | 31 | **32** | **33** | 34 | 35 | 36 |
|  | ***16*** | 37 | 38 | 39 | 40 | 41 | 42 | 43 | 44 | 45 | 46 | 47 | 48 |
|  | ***17*** | 49 | 50 | 51 | 52 | 53 | 54 | 55 | 56 | 57 | 58 | 59 | 60 |
|  | ***18*** | 61 | 62 | 63 | 64 | 65 | 66 | 67 | 68 | 69 | 70 | 71 | 72 |
|  | ***19*** | 73 | 74 | 75 | 76 | 77 | 78 | 79 | 80 | 81 | 82 | 83 | 84 |
|  | ***20*** | 85 | 86 | 87 | 88 | 89 | 90 | 91 | 92 | 93 | 94 | 95 | 96 |
|  | ***21*** | 97 | 98 | 99 | 100 | 101 | 102 | 103 | 104 | 105 | 106 | 107 | 108 |
|  | ***22*** | 109 | 110 | 111 | 112 | 113 | 114 | 115 | 116 | 117 | 118 | 119 | 120 |
|  | ***23*** | 121 | 122 | 123 | 124 | 125 | **126** | 127 | **128** | **129** | 130 | 131 | 132 |
|  | ***24*** | 133 | 134 | 135 | 136 | 137 | 138 | 139 | 140 | 141 | 142 | 143 | 144 |
|  | ***25*** | 145 | 146 | 147 | 148 | 149 | 150 | 151 | 152 | 153 |  |  |  |

Pep#32

Pep#30

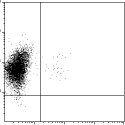

1.5%

Pep#126	Pep#33

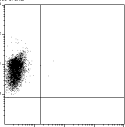

0.1%

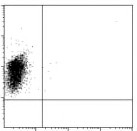

0.1%

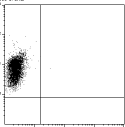

0.1%

Pep#128	Pep#129

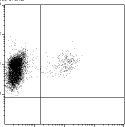

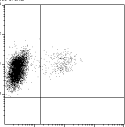

4.4%

4.3%

IFNγ

CD3

**Supp. figure 4. MSLN epitope identification.** (A) To identify immunogenic peptide epitopes T cell lines were screened against individual MSLN minipools by IFNγ ELISpot. Panel A shows results for donor 11 (results reported as SFC/2×10^5^), with the potential stimulatory peptides shown in panel B. (C) To identify the immunogenic epitopes the MSLN T cell line was exposed to each of the peptides identified in panel B – screening was performed by ICS for IFNγ and results reported as % of CD3+/IFNγ+ cells.

A Control

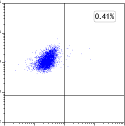

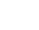

0.4%

CD4

Donor 3

Pep #72

CD4

B Control

Donor 9

Pep #72

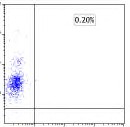

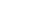

0.2%

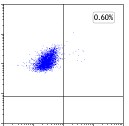

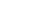

0.6%

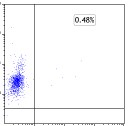

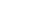

0.4%

Donor 5

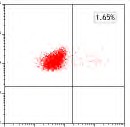

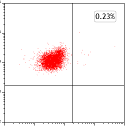

0.2%

1.7%

0

IFNγ

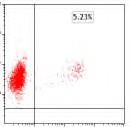

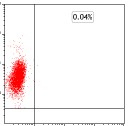

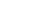

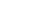

0.1%

5.2%

IFNγ

CD8

CD8

C Control	Pep #72

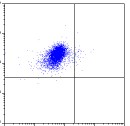

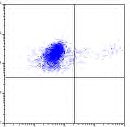

**D**

700

Donor 5

0.2%

3.2%

0

CD4

600

500

SFC /2x10^5^ cells

400

300

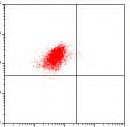

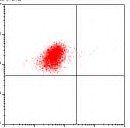

0.3%

0.4%

IFNγ

200

100

0

NO ab ABC	DR	DQ	DP

Pep #72

CD8

**Supp. figure 5. Detection of CD4- and CD8-mediated reactivity to peptide #72.** (A-C) ICS (IFNγ production) analysis to determine whether reactivity to peptide #72 was detected in the CD4+ or CD8+ T cell compartment for donors 3 (A), 9 (B) and 5 (C). For donor 5, the observed CD4-mediated reactivity was further interrogated by IFNγ ELISpot (D) in the absence or presence of MHC class I or class II blocking antibodies. Results are reported as SFC/2x10^5^ input cells.

A Control	Pep #132

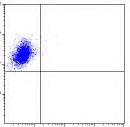

0.3%

Donor 1

CD4

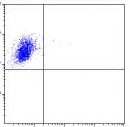

0.3%

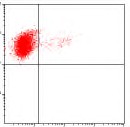

5.2%

IFNγ

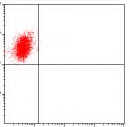

0.1%

CD8

Pep #132

40

35

30

25

% Lysis

20

15

10

5

0

auto A2 A11 B35 B38

HLA-A HLA-B

B Control

IFNγ

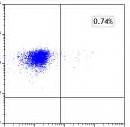

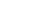

0.7%

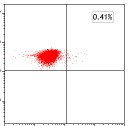

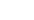

0.4%

E:T 40:1

Donor 7

CD4

CD8

Pep #151

Pep #151

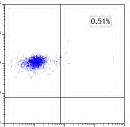

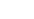

0.5%

60

50

40

E:T 40:1

% Lysis

30

20

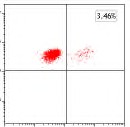

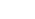

3.5%

10

0

auto A2 A25 B15 B51

HLA-A HLA-B

C Control	Pep #34

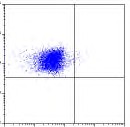

0.4%

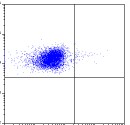

0.5%

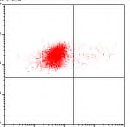

2.2%

Donor 5

CD4

IFNγ

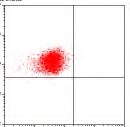

0.2%

CD8

**D**

Pep #34

35

30

E:T 40:1

25

% Lysis

Donor 3

CD4

20

15

10

5

0

CD8

auto A2 A24 B13 B35

HLA-A HLA-B

Control	Pep #118

_50_	Pep #118

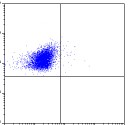

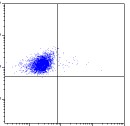

1.2%

0.8%

40

E:T 40:1

30

% Lysis

20

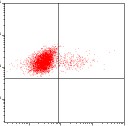

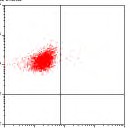

0.5%

4.8%

IFNγ

10

0

auto A11 A24 B35 B40

HLA-A HLA-B

**Supp. Figure 6. Identification of HLA-restricting alleles for the immunodominant MSLN peptides.** (A) For peptide 132: ICS (IFNγ production) analysis to determine whether reactive T cells were detected in the CD4+ or CD8+ T cell compartment (left panel). ⁵¹Cr release assay (5-8 hr co-culture) using peptide-pulsed autologous and partially HLA-matched PHA blasts as targets (right panel). Results show % specific lysis (mean±SD), E:T 40:1. Panels B-D show similar data for peptides 151, 34, and 118, respectively.

**A**

800

**B**

900

| **Pep#145** | |
| --- | --- |
| **Pep #145** | **GLQGGIPNGYLVLDL** |
| **9mers** | **GLQGGIPNG** |
|  | **LQGGIPNGY** |
|  | **QGGIPNGYL** |
|  | **GGIPNGYLV** |
|  | **GIPNGYLVL** |
|  | **IPNGYLVLD** |
|  | **PNGYLVLDL** |

600

SFC/2x10^5^ cells

SFC/2x10^5^ cells

| **Pep#30** | |
| --- | --- |
| **Pep #30** | **ALPLDLLLFLNPDAF** |
| **9mers** | **ALPLDLLLF** |
|  | **LPLDLLLFL** |
|  | **PLDLLLFLN** |
|  | **LDLLLFLNP** |
|  | **DLLLFLNPD** |
|  | **LLLFLNPDA** |
|  | **LLFLNPDAF** |

600

400

300

200

0

30	ALP	LPL	PDL	LDL	DLL	LLL	LLF	neg

0

145 GLQ LQG QGG GGI GIP IPN PNG neg

pep

9-mer

ctrl

pep

9-mer

ctrl

**C	D**

| **Pep #91** | |
| --- | --- |
| **Pep #91** | **PQGYPESVIQHLGYL** |
| **14mer 13mer 12mer 11mer 14mer 13mer 12mer**  **11mer** | **QGYPESVIQHLGYL GYPESVIQHLGYL YPESVIQHLGYL PESVIQHLGYL**  **PQGYPESVIQHLGY PQGYPESVIQHLG PQGYPESVIQHL**  **PQGYPESVIQH** |
| **10mers** | **PQGYPESVIQ QGYPESVIQH**  **GYPESVIQHL YPESVIQHLG**  **PESVIQHLGY**  **ESVIQHLGYL** |
| **9mers** | **PQGYPESVI QGYPESVIQ**  **GYPESVIQH YPESVIQHL**  **PESVIQHLG ESVIQHLGY**  **SVIQHLGYL** |

| **Pep#128/129** | |
| --- | --- |
| **Pep #128** | **KALSQQNVSMDLATF** |
| **Pep #129** | **QQNVSMDLATFMKLR** |
| **9mers** | **KALSQQNVS** |
|  | **ALSQQNVSM** |
|  | **LSQQNVSMD** |
|  | **SQQNVSMDL** |
|  | **QQNVSMDLA** |
|  | **QNVSMDLAT** |
|  | **NVSMDLATF** |
|  | **VSMDLATFM** |
|  | **SMDLATFMK** |
|  | **MDLATFMKL** |
|  | **DLATFMKLR** |

700	90

600	^80^

70

SFC/2x10^5^ cells

SFC/2x10^5^ cells

500	_60_

400	50

300	40

200	^30^

20

100	_10_

0	0

Pep

91

QGY GYP YPE PES PQG PQG PQG PQG PQG QGY GYP YPE PES ESV PQG QGY GYP YPE PES ESV SVI

neg

128

129

KAL ALS LSQ SQQ QQN QNV NVS VSM SMD MDL DLA

neg

14 13 12 11 14 13 12 11	10	9

Pep

Pep

pep

-mer

ctrl

pep

9-mer

ctrl

**Supp. figure 7. Identification of minimal epitopes for the immunodominant MSLN peptides.** (A) For peptide 30: the 9mer LPLDLLLFL was identified as the minimal recognized epitope using an array of overlapping 9mers for stimulation of MSLN-STs (top panel) and the IFNγ ELISpot as a readout (bottom panel). Panels B-D show similar data for peptides 145, 91, and 128/129, respectively. Results are reported as SFC/2x10^5^ input cells.
